# Hippocampal Subfield Volumes in Abstinent Men and Women with a History of Alcohol Use Disorder

**DOI:** 10.1101/715292

**Authors:** Kayle S. Sawyer, Noor Adra, Daniel M. Salz, Maaria I. Kemppainen, Susan M. Ruiz, Gordon J. Harris, Marlene Oscar-Berman

**Affiliations:** VA Boston Healthcare System, Boston, MA 02130; Boston University School of Medicine, Boston, MA 02118; Massachusetts General Hospital, Boston, MA 02129; Sawyer Scientific, LLC, Boston, MA 02120; Harvard Medical School, Boston, MA 02115

**Keywords:** Alcohol, hippocampal subfields, gender, memory, brain, alcohol use disorder

## Abstract

Alcohol use disorder (AUD) has been associated with abnormalities in hippocampal volumes, but these relationships have not been fully explored with respect to sub-regional volumes, nor in association with individual characteristics such as gender differences, age, and memory. The present study examined the impact of those variables in relation to hippocampal subfield volumes in abstinent men and women with a history of AUD. Using Magnetic Resonance Imaging at 3 Tesla, we obtained brain images from 67 participants (31 women) with AUD and 63 healthy control (NC) participants (30 women) without AUD. We used Freesurfer 6.0 to segment the hippocampus into 12 regions. These were imputed into mixed models to examine the relationships of brain volume with AUD group, gender, age, drinking history, and memory. The AUD group had approximately 5% smaller CA1, hippocampal tail, and molecular layer regions than the NC group. Age was negatively associated with volumes for the AUD group in the hippocampal tail, subiculum, and presubiculum. The relationships for delayed and immediate memory with hippocampal tail volume differed for AUD and NC groups: Higher scores were associated with smaller volumes in the AUD group, but larger volumes in the NC group. Length of sobriety was associated with decreasing CA1 volume in women (0.02% per year) and increasing volume size in men (0.03% per year). These findings confirm and extend evidence that AUD, gender, age, and abstinence differentially impact volumes of component parts of the hippocampus. The course of abstinence on CA1 volume differed for men and women, and the differential relationships of subregional volumes to age and memory could indicate a distinction in the impact of AUD on functions of the hippocampal tail.

## Introduction

Magnetic resonance imaging (MRI) has been used extensively to study morphological changes in the brain associated with alcohol use disorder (AUD), a widespread and harmful condition [1, 2]. Because memory impairments are associated with long-term chronic AUD, one neuroanatomical focus of investigation has been the hippocampus [3]. Moreover, findings from previous research have shown that the largest subcortical volume loss observed in the brains of people with chronic AUD was in the hippocampus [4], and a meta-analysis [5] found a negative association between total hippocampal volume and degree of alcohol use (clinical vs. subclinical).

Although results of MRI studies have suggested gender-specific susceptibility of the brain to alcohol abuse [6–15], few studies have examined gender differences in the relationship between hippocampal volume and AUD. Instead, researchers have assessed AUD-related hippocampal abnormalities in men [16–20], or in AUD groups comprised of a combination of men and women [21–23]. In studies that did analyze men with AUD (AUDm) and women with AUD (AUDw) separately, investigators observed lower hippocampal volumes in the AUDm and in the AUDw, but they did not find significant interactions between gender and diagnostic group [21, 23], while we previously observed smaller hippocampal volumes for AUDm than AUDw [9].

Similarly, the relationships observed between hippocampal volume and cognitive ability also have been inconsistent. It has been widely reported that lesions to the hippocampus result in memory deficits [24, 25]. Positive correlations between hippocampal volume and memory performance also have been reported in clinical populations such as patients with Alzheimer’s disease or amnesic disorders [26–29], but this positive correlation has not been systematically observed in healthy populations [30]. In the context of AUD, some research has indicated the presence of functional deficits with lower hippocampal volumes. For example, decreases in hippocampal gray matter volume in alcohol dependent men have been associated with executive functioning deficits [18] as measured by the Trail Making Test Part B, an assessment of visual-conceptual and visual-motor tracking skills [31]. However, among participants with AUD in these earlier studies, smaller volumes of the anterior portion of the hippocampus were not significantly associated with memory impairment [32].

The inconsistent findings relating volume and function may be impacted by the distinct functions of hippocampal subfields and the processing streams that exist within the hippocampus (Figure 1, which overlays this processing stream on an MRI atlas; [33]). Traditionally, based upon research involving anatomical dissection, histology, and physiology, hippocampal subfields have been defined as the subiculum, dentate gyrus, and *cornu ammonis* regions (CA1 through CA3) [34]. Researchers have focused on two major neural pathways, as follows: First, major input to the hippocampus originates in entorhinal cortex and proceeds along a trisynaptic circuit (1) to the dentate gyrus, then (2) to CA2 or CA3, and finally, (3) to CA1 (and then, subiculum), which serve as outputs of the hippocampus. Second, the entorhinal cortex (not part of hippocampus) additionally has a direct connection to CA1, among other regions. All the input connections from CA1, CA2, CA3, and subiculum are made within an area called the molecular layer (also known as *stratum lacunosum moleculare* or SLM).

**Figure 1.**
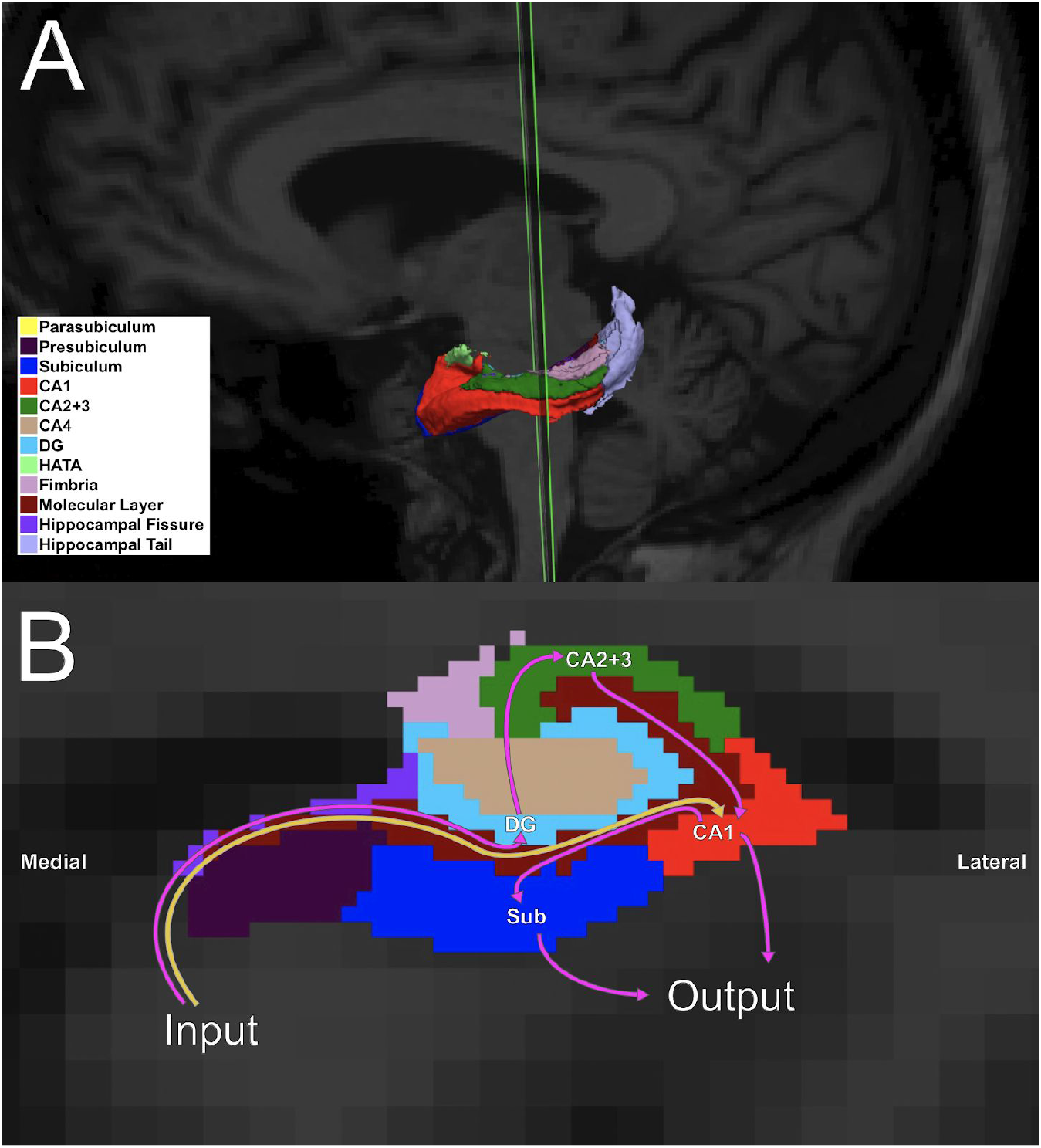
Hippocampal subfield segmentation using the procedure by Iglesias and colleagues [33]. A. This three-dimensional model of the left hippocampus depicts its segmentation into 12 subfields (although not all subfields are visible from the angle presented). B. This coronal section through the hippocampus illustrates subfields that are visible in the slice, and it shows the two major hippocampal neural pathways originating in the entorhinal cortex and as described in the text: The pink arrows indicate the trisynaptic circuit; the yellow arrow shows the direct circuit. Although the connection from CA1 to the subiculum is not explicitly part of the trisynaptic circuit, it is important as an output region, in addition to CA1. Abbreviations: CA1 = cornu ammonis 1; CA2+3 = cornu ammonis 2 and 3; CA4 = cornu ammonis 4; DG = dentate gyrus; HATA = hippocampal-amygdaloid transition area; Sub = subiculum.

The molecular layer is the point of entry for information, including the direct pathway from entorhinal cortex that is theorized to contain information regarding presently experienced stimuli [35, 36]. The dentate gyrus has been shown to play a role in distinguishing different contexts [37]. For the structures involved in the trisynaptic pathway, the CA2 and CA3 regions have been ascribed to learning, encoding [38], early retrieval of verbal information [39], and disambiguation and encoding of overlapping representations [40]. The output fields, CA1 and subiculum, are thought to compare current context with remembered contexts [41]. Their projections include regions implicated in addictive disorders: the prefrontal cortex, amygdala, nucleus accumbens, and ventral tegmental area. This processing stream is repeated along the axis of the hippocampus, described in the human as anterior to posterior, and in rodent models as ventral to dorsal. The anterior (or ventral) portion has been considered to involve a wide variety of contexts including stress, emotion, and affect [42], while the posterior (or dorsal) portion has been demonstrated to have involvement in spatial processing. White matter fibers that carry the projections of the hippocampus include the fimbria [43].

In a sample of men with ongoing AUD, Lee et al. [17], reported smaller volumes of the left presubiculum, bilateral subiculum regions, and fimbria, but in studies that have included women, significant differences in interactions of AUD and gender have not been detected for hippocampal subfield volumes [22, 23], perhaps due to insufficient sample sizes. Research also has supported reports of smaller AUD-related volumes in subiculum, CA1, CA4, or other parts of the hippocampus (dentate gyrus, hippocampal-amygdaloid transition area [HATA], and the fimbria) [17, 23], along with CA2 and CA3, which showed significant normalization two weeks following withdrawal [22]. Additionally, effects of age were found for the fimbria and the hippocampal fissure, while a significant interaction between AUD diagnosis and age was seen in CA2 and CA3 regions. However, in neither the Lee et al.[17] nor the Kühn et al. [22] studies, was memory assessed, and Zahr et al. [23] found no significant memory impairment associated with reduced hippocampal subfield volumes.

In summary, the present study examined the impact of gender on hippocampal subfield volumes, and the relationships between age, memory, and volumes of these regions in abstinent individuals with a history of AUD. Examining the impact of AUD on individual subfield volumes and their interactions with age, gender, and memory can further elucidate which components of this brain network are associated with the problems contributing to, and resulting from, AUD.

## Methods

As described in detail below, 67 AUD (31 women) and 63 NC participants (30 women) were included in analyses (Table 1). Memory and drinking history were assessed. T1-weighted 3T MRI scans were obtained with 1×1×1.5 mm voxels. The 12 hippocampal subfield volumes were obtained with FreeSurfer 6.0. In the mixed models, volume was entered as the dependent variable, and ‘group,’ ‘gender,’ ‘region,’ and ‘age’ were the independent factors. We examined the interaction of group-by-gender-by-region to assess how the impact of AUD differed for men and women. Also, we examined the interaction of group-by-gender-by-age to assess abnormal aging for men as compared to women. Following significant effects, additional models were constructed to examine relationships with memory measures, and with drinking history.

**Table 1.** Participant Characteristics. Minimum (Min), Maximum (Max), Means, and standard deviations (SD) are provided for the participant characteristics of AUDw (N=31) and AUDm (N=36) (women and men with a history of Alcohol Use Disorder), along with NCw (N=30) and NCm (N=33) (women and men without a history of AUD). DMI was not obtained from one AUDm and one NCm. IMI was not obtained from one AUDm and one NCm. Abbreviations: FS = Wechsler Adult Intelligence Full Scale IQ; DMI = Delayed Memory; IMI = Immediate Memory; DHD = Duration of Heavy Drinking; DD = Daily Drinks; LOS = Length of Sobriety.

| Measure | AUDw |  |  |  | AUDm |  |  |  |
| --- | --- | --- | --- | --- | --- | --- | --- | --- |
|  | Mean | SD | Min | Max | Mean | SD | Min | Max |
| Age (years) | 54.5 | 12.0 | 31.1 | 76.0 | 51.2 | 10.8 | 28.1 | 76.8 |
| Education (years) | 14.7 | 1.8 | 12.0 | 19.0 | 14.1 | 1.9 | 12.0 | 18.0 |
| FS | 103.1 | 14.5 | 69.0 | 132.0 | 104.1 | 16.6 | 75.0 | 142.0 |
| IMI | 107.3 | 17.3 | 56.0 | 135.0 | 102.6 | 14.9 | 73.0 | 132.0 |
| DMI | 110.2 | 16.9 | 62.0 | 135.0 | 104.8 | 14.1 | 81.0 | 133.0 |
| DHD (years) | 13.7 | 5.6 | 5.0 | 25.0 | 17.5 | 9.1 | 5.0 | 37.0 |
| DD (oz etOH/day) | 8.8 | 7.2 | 2.5 | 34.8 | 11.8 | 6.8 | 2.3 | 26.6 |
| LOS (years) | 10.9 | 11.9 | 0.1 | 36.1 | 3.9 | 6.2 | 0.1 | 27.4 |

**Table 1. Participant Characteristics.**
| Measure | NCw |  |  |  | NCm |  |  |  |
| --- | --- | --- | --- | --- | --- | --- | --- | --- |
|  | Mean | SD | Min | Max | Mean | SD | Min | Max |
| Age (years) | 53.0 | 15.8 | 27.8 | 78.4 | 50.5 | 12.3 | 26.6 | 81.7 |
| Education (years) | 15.3 | 2.6 | 12.0 | 20.0 | 15.4 | 2.5 | 10.0 | 20.0 |
| FS | 110.8 | 15.1 | 82.0 | 141.0 | 112.7 | 14.7 | 89.0 | 141.0 |
| IMI | 113.0 | 16.1 | 89.0 | 143.0 | 110.5 | 15.4 | 84.0 | 140.0 |
| DMI | 115.0 | 12.5 | 93.0 | 146.0 | 114.2 | 17.4 | 78.0 | 147.0 |
| DHD (years) | 0.0 | 0.0 | 0.0 | 0.0 | 0.0 | 0.0 | 0.0 | 0.0 |
| DD (oz etOH/day) | 0.3 | 0.3 | 0.0 | 1.1 | 0.2 | 0.2 | 0.0 | 0.7 |
| LOS (years) | 2.2 | 7.4 | 0.0 | 31.9 | 2.1 | 7.9 | 0.0 | 40.2 |

### Participants

The study originally included 146 participants: 73 abstinent adults with a history of chronic AUD (33 AUDw; 40 AUDm) and 73 adult controls without AUD (34 NCw; 39 NCm), who were recruited locally through online and print advertisements. Participants provided written informed consent for participation in the study, which was approved by the Institutional Review Boards at the Boston VA Healthcare System, Massachusetts General Hospital, and Boston University School of Medicine. Exclusion criteria for participants included left-handedness, Korsakoff’s syndrome, HIV, hepatic disease, head injury with loss of consciousness greater than 15 minutes, stroke, seizures unrelated to AUD, schizophrenia, Hamilton Rating Scale for Depression (HRSD) [44] score over 14, and illicit drug use (except marijuana) greater than once a week within the past five years. Sixteen individuals were excluded from the study for the following reasons: Three AUD participants (1 woman) were excluded for illicit drug use, and another three (1 woman) for brain lesions or head trauma. Six NCm were excluded for binge drinking, head trauma, or unusable scan data. Two NCw were excluded for claustrophobia or brain lesions, and another two NCw were identified as outliers, with brain hippocampal region volumes +/-4 standard deviations, and removed from further analyses. The final sample of participants was comprised of a total of 67 AUD (31 women) and 63 NC participants (30 women) included in data analyses (Table 1).

Diagnostic criteria for study exclusion were based on a medical history interview, the HRSD, and a computer-assisted, shortened version of the Computerized Diagnostic Interview Schedule (DIS) [45]. The DIS provides diagnoses of lifetime psychiatric illnesses as defined by criteria established by the American Psychiatric Association. Full-Scale IQ and cognitive performance were measured through the Wechsler Adult Intelligence Scale-IV (WAIS-IV) and the Wechsler Memory Scale-IV (WMS-IV) [46]. The memory assessments were conducted by trained researchers and consisted of tasks designed to measure memory for short stories, lists of words, shapes, and spatial locations. Incomplete WMS-IV scores were obtained from one NCm and two AUDm and were excluded from the memory analyses.

A number of participants were taking medications for a variety of conditions, had used drugs earlier than five years before enrollment, or had a potentially confounding medical history. We included these participants with confounding factors, so that our sample would be more representative of the conditions present in the United States, thereby allowing for greater generalizability of the results. However, the presence of confounding factors in the sample may limit the interpretability of our findings. Therefore, in the analysis of the results, a subsample of 77 participants (26 AUD; 51 NC) was created consisting of “unconfounded” participants who were not currently taking psychotropic medications, and reported never using illicit drugs more than once a week. Additionally, that subsample was restricted to individuals for whom no source indicated hepatitis or cirrhosis, nor any of the following disorders: major depressive, bipolar I or II, schizoaffective, schizophreniform, or generalized anxiety. All statistical effects of group reported in this manuscript (including group interactions) remained significant for this unconfounded subsample.

Drinking history was assessed using Duration of Heavy Drinking (DHD), i.e., years of consumption of 21 drinks or more per week, and Length of Sobriety (LOS), which measures abstinence duration in years. The amount, type, and frequency (ounces of ethanol per day, roughly corresponding to daily drinks; DD) of alcohol use was measured for the last six months during which the participant drank alcohol [47]. Criteria for AUD participants included at least five years of previous alcohol abuse or dependence, and a minimum of four weeks of abstinence prior to testing. Two NCw and one NCm had no prior history of drinking, whereas the remaining NC participants drank occasionally. Compared to the AUDw, the AUDm had greater periods of heavy drinking and shorter periods of abstinence, which are consistent with national trends [48] and allow for generalizability of the results. However, to improve interpretability, we created a subsample in which the AUDw and AUDm were not significantly different by removing four AUDw with the longest LOS and shortest DHD values. All statistical effects of gender reported in this manuscript (including gender interactions) remained significant for this subsample.

### MRI Acquisition and Analysis

MRI scans were obtained at the Martinos Center for Biomedical Imaging at Massachusetts General Hospital on a 3 Tesla Siemens (Erlangen, Germany) MAGNETOM Trio Tim scanner with a 32-channel head coil. Image acquisitions included two T1-weighted multiecho MPRAGE scans collected for volumetric analysis that were averaged to aid in motion correction (TR = 2530 ms, TE = 1.79 ms, 3.71 ms, 5.63 ms, 7.55 ms [root mean square average used], flip angle = 7 degrees, field of view = 256 mm, matrix = 256 × 256, slice thickness = 1 mm with 50% distance factor, 176 interleaved sagittal slices, GRAPPA acceleration factor = 2; voxel size = 1.0 mm × 1.0 mm × 1.5 mm).

Scans were analyzed using an automated hippocampal segmentation method [33] in FreeSurfer (https://surfer.nmr.mgh.harvard.edu), which more recently has shown high reliability and agreement with manual segmentations [49]. Brain reconstructions were manually inspected and errors were corrected. Volumes of the hippocampal subfields (12 per hemisphere) were calculated using the hippocampal subfields subroutines for Freesurfer 6.0 (which omits the alveus due the poor reliability of the segmentation). These 12 volumes were used to define the extent of the hippocampus. Estimated total intracranial volume was taken from the segmentation volume estimate [50]. The volumes were divided by estimated total intracranial volume (eTIV) to adjust for head size.

### Statistical Analyses

All statistical analyses were performed using R version 3.4.0 [51]. We used hierarchical linear models [52] to investigate the impact of several variables on regional hippocampal volumes. Data and code are available at https://gitlab.com/kslays/moblab-hippocampus. Because brain volumes vary with head size, we used normalized volume values (i.e., volume of each participant divided by total intracranial volume). Left and right hemisphere values of corresponding regions were averaged. Participants with outlier volumes (greater than or less than four standard deviations) within any region were removed from the analysis. In the regression models, we visually confirmed that other regression assumptions were satisfied (normality, equality of variance, homogeneity of regression), and set thresholds for multicollinearity (Pearson correlations among predictors were < 0.5) and influence (Cook’s D < 1.0).

We first conducted an analysis of total hippocampal volume using multiple regression. We constructed a model predicting whole hippocampal volume from the interaction of ‘group’, ‘gender’, and ‘age’, along with the lower-order interactions and main effects. Next, for our analysis of subfield volumes, we used mixed models. Volume was entered as the dependent variable, and ‘group,’ ‘gender,’ ‘region,’ and ‘age’ were the independent factors (all fixed effects). In order to account for multiple observations per subject (i.e., volumes for each ‘region’), individual subject effects were specified as random intercepts.

Interactions were included to assess the primary aims of this project. We examined the interaction of group-by-gender-by-region to assess how the impact of AUD differed for men and women, and how gender differences impacted certain regions in comparison to others. Also, we examined the interaction of group-by-gender-by-age to assess abnormal aging for men as compared to women. The four way interaction of group-by-gender-by-region-by-age was examined to confirm homogeneity of regression slopes. All the non-significant (i.e., *p* > 0.05) interactions (except lower-order interactions included in higher-order interactions) that had been added to confirm homogeneity of regression slopes were removed, and the subsequent model was used to report results.

Following significant interactions, we evaluated mean differences within each region using Bonferroni multiple comparisons correction of a family-wise *p*-value threshold of 0.05. There were 12 hippocampal subfields, so this correction resulted in an adjusted *p*-value threshold of 0.0042. Results were reported as percent differences (i.e., mean difference/mean). Significant regions were then investigated for our secondary aims, i.e., analyses involving memory scores and drinking history. For interactions of group-by-gender, the comparisons of AUDm vs. NCm and AUDw vs. NCw were planned, while for region-by-group, only AUD vs. NC comparisons were planned. For age interactions, the effect of age for each of the subgroups was assessed, and the slope differences were compared in the same manner as mean differences.

To address our secondary aims, we started with the model that resulted from our primary aims. We then investigated interactions of group-by-gender with separate models for two memory measures from the WMS-IV: the immediate memory index (IMI) and the delayed memory index (DMI). These interactions were assessed in conjunction with each score’s interaction with each region. We applied the same procedure used in the primary analyses to examine heterogeneity of regression slopes and to remove extraneous interactions. Following significant interactions, we examined the significance of each score’s slope in the same 12 regions as for the primary analyses, and the same slope differences as for the primary analyses. The WMS-IV Index Scores and WAIS-IV Composite Scores were analyzed in separate ANOVA models, each of which contained between-subjects factors of group and gender, and a group-by-gender interaction term.

To investigate how alcohol-drinking history (DD, DHD, and LOS) impacted volume, we conducted an analysis with only the AUD participants included. The NC participants were not included because the participants did not have meaningful variation within the scores. We examined interactions of each of these three measures with gender-by-region in a single model. As with the primary and secondary aims, we tested for heterogeneity of regression slopes, and removed interactions as dictated by significance levels. Following significant interactions, we examined the significance of each score’s slope in the same 12 regions as for the primary analyses, and we compared the slopes for men to the slopes for women.

## Results

### Participant Characteristics

Table 1 provides information about the participants. The groups consisted of 67 AUD (31 women) and 63 NC (30 women), with a mean age of 52 years for the 130 participants. Between-group (NC vs. AUD) differences in age were not significant (95% CI of mean difference = [5.4, −3.4] years). The AUD participants had 0.9 fewer years of education (*t*(114.74) = −2.49, *p* < 0.05) and 8.1 points lower FSIQ scores (*t*(127.99) = −3.07, *p* < 0.01). The AUDw had significantly longer LOS and DHD than the AUDm; other measures did not differ significantly. However, as described in the Methods, all gender effects reported remained significant in a subsample for which the AUDw and AUDm did not differ significantly by LOS or DHD.

### Hippocampus Volumes, Group, Gender, and Age

All volumes were converted to proportions of head size (eTIV) before analysis. Linear regression analyses of whole hippocampus volume revealed a significant interaction of group-by-age (F(1, 122) = 7.15, *p* < 0.01). The AUD group had 5.14% smaller volumes than the NC group (t(122) = −3.855, *p* < 0.001). Age was associated with a 0.44%/year decline for the AUD group, which was significantly steeper than the 0.13%/year decline observed for the NC group.

To analyze the subfield volumes, we used mixed-model regression. These analyses revealed significant three-way interactions (Table S2) for group-by-region-by-age (F(11, 5366.83) = 3.02, *p* < 0.01) and gender-by-region-by-age (F(11, 5366.83) = 2.50, *p* < 0.01). No significant group-by-gender interactions were observed. Group differences in means, and in slopes (for age), are reported below.

Mean differences between groups revealed significantly (Bonferroni adjustment of family-wise *p* < 0.05 for 12 regions: *p* < 0.0042) smaller CA1, hippocampal tail, and molecular layer volumes in the AUD group compared to the NC group, with marginal significance (*p* = 0.0046) for subiculum (Figure 2 and Table S2). The AUD group had 5.18%, 5.08%, 4.80%, and 3.70% smaller volumes in the CA1, molecular layer, hippocampal tail regions, and subiculum, respectively.

**Figure 2.**
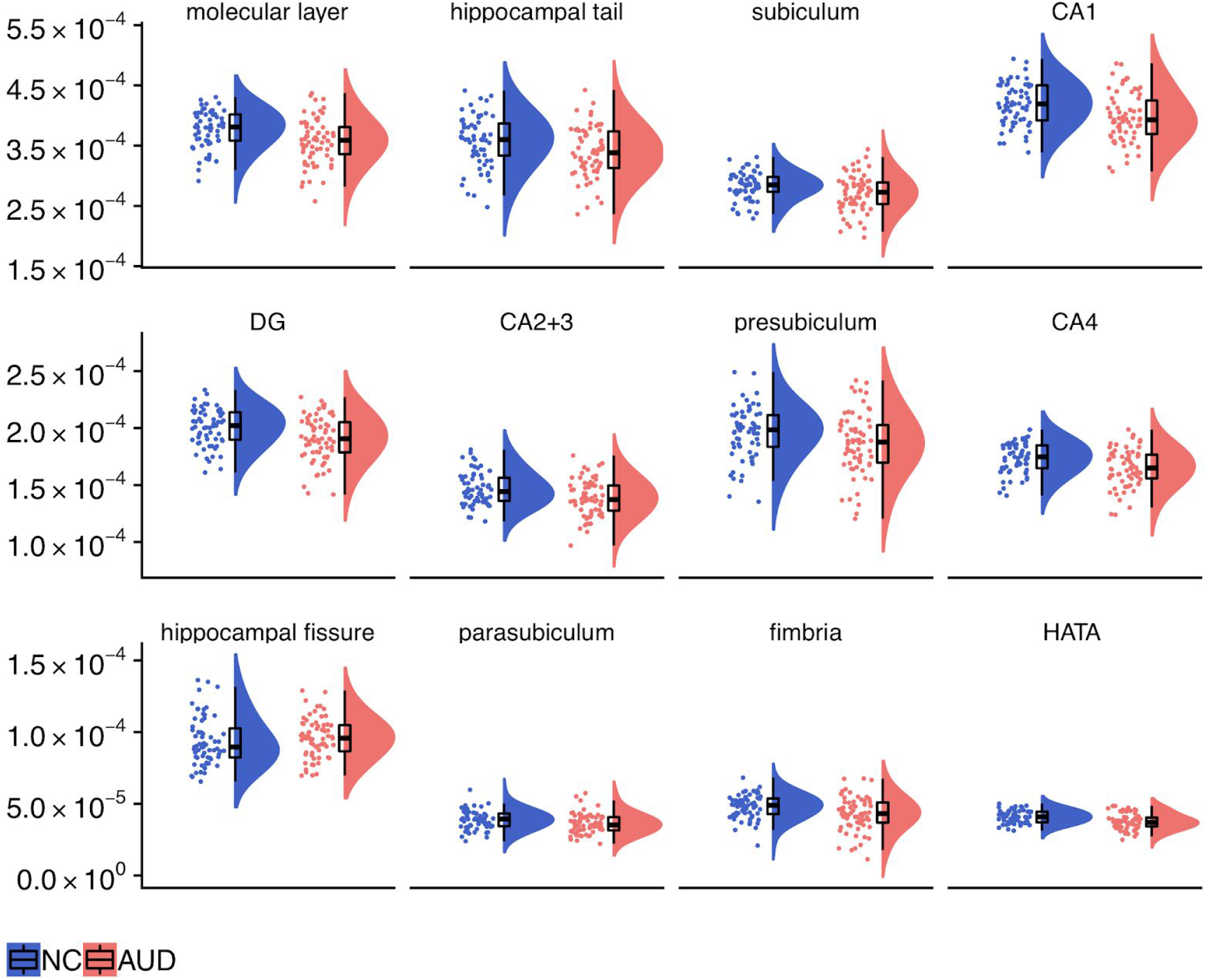
Regional volumes (proportion of estimated total intracranial volume). Half violin raincloud plots [53] show hippocampal subfield volumes for the NC and AUD groups. See Table S1 for mean and standard deviation values. Abbreviations: CA1 = cornu ammonis 1; CA2+3 = cornu ammonis 2 and 3; CA4 = cornu ammonis 4; DG = dentate gyrus; HATA = hippocampal-amygdaloid transition area; Sub = subiculum. *Indicates regions where AUD < NC, *p* < 0.0042.

In the entire sample of the 130 participants, significant gender differences (with analyses including AUD and NC participants together) were found for the subiculum, presubiculum, CA1, and molecular layer regions. Women had 8.43%, 4.99%, 4.47%, and 3.30% larger volumes in the presubiculum, subiculum, molecular layer, and CA1, respectively. As with all analyses, volumes had been first converted to a proportion of cranial volume.

The impact of age on subfield volumes interacted significantly with group. For the AUD group, age was associated with significantly smaller hippocampal tail, subiculum, presubiculum, and molecular layer volumes (−0.52, −0.45, −0.58, and −0.47, respectively; all percent/year); these relationships were significantly more negative than those observed for the NC group (−0.17,-0.03, −0.04, and −0.17; all percent/year). The impact of age on subfield volumes also significantly interacted with gender. For women, age was associated with 0.4 percent/year lower CA1 volumes, a rate significantly less negative than that observed for men (1.0 percent/year).

### Subfield Volumes and Memory

Following our main analyses, we assessed the relationship of memory scores and volume. We found significant interactions (Figure 3, Figure S1, Figure S2, Table S3, and Table S4) for group-by-region-by-IMI and for group-by-region-by-DMI. For better interpretability, slopes were divided by the grand mean for each region and are thus presented as percent volume/IMI or DMI unit.

**Figure 3.**
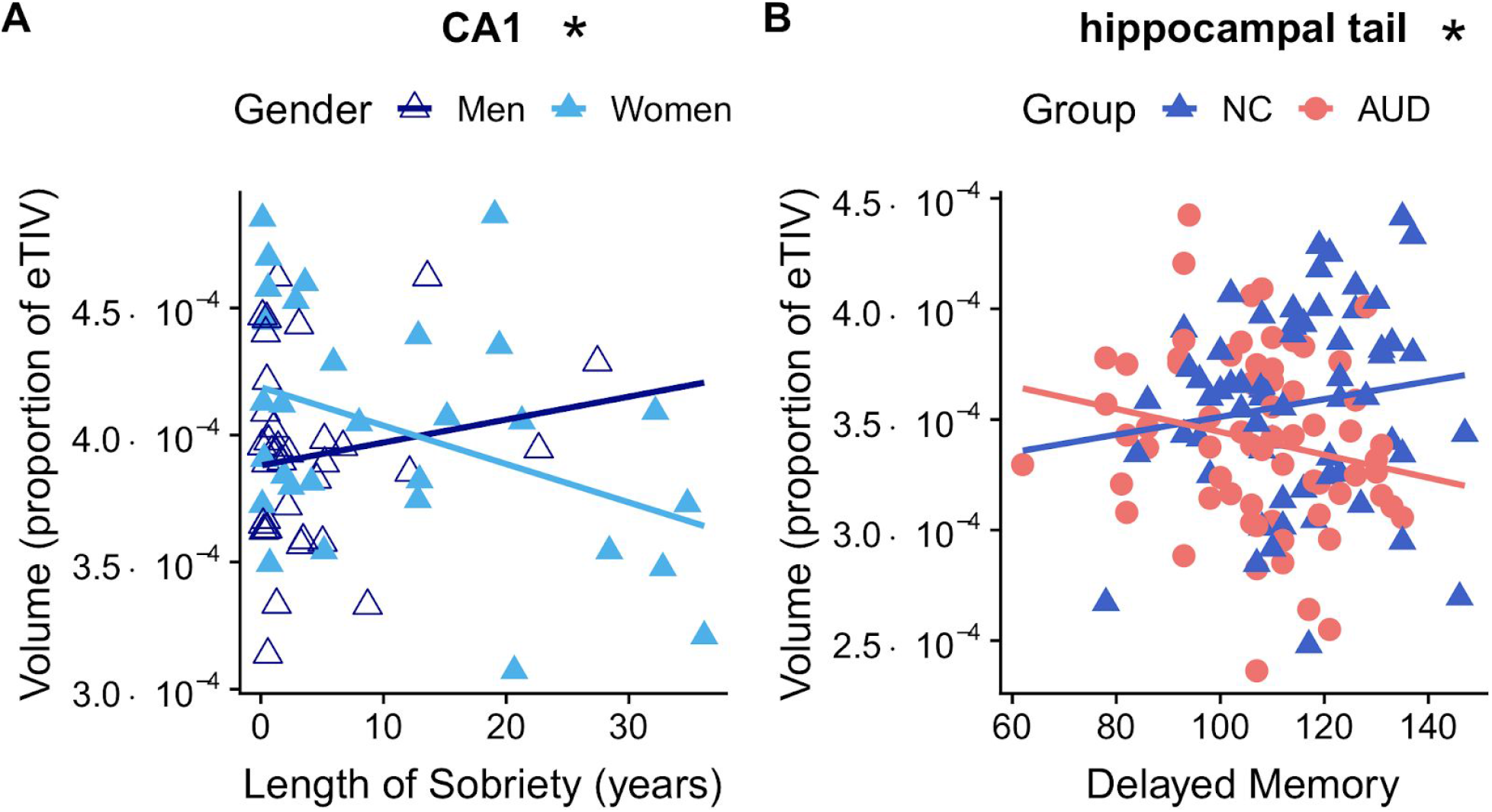
Distinct relationships of subfield volume for AUD and NC groups. A. For AUD men, CA1 volumes (proportion of eTIV) were positively associated with Length of Sobriety (years), while for AUD women, a negative relationship was observed. B. For the AUD group, Delayed Memory was associated with lower hippocampal tail volumes, while for the NC group, a positive relationship was observed. Abbreviations: CA1 = cornu ammonis 1; eTIV = estimated total intracranial volume * Indicates regions with interactions significant at *p* < 0.0042.

#### Immediate Memory Index

Following the identification of significant group effects for CA1, hippocampal tail, and molecular layer, and of marginal significant group effects in the subiculum (*p* = 0.0046), we found that associations of IMI with those regions differed for AUD and NC groups in the hippocampal tail (t(594.84) = −3.51, *p* < 0.001). In AUD participants, the volumes decreased by 0.12% per unit of IMI, while for NC they increased by 0.13% with each unit of IMI.

#### Delayed Memory Index

Also in the hippocampal tail, a significant interaction of AUD and NC participants was observed between DMI and subfield regions (Table S4). In AUD participants, the volumes were 0.14% lower with each unit of DMI, while for NC participants, volumes were 0.20% higher with each unit of DMI, relationships that differed significantly from each other (t(599.16) = −4.55, *p* < 0.0001).

### Subfield Volumes and Drinking History

For the AUD group, we examined the relationships of DHD, DD, and LOS to volumes, and the interactions of those drinking history measures. All models included age as a covariate, and correlations between age, DHD, DD, and LOS were low (all r < 0.5; age with DHD = 0.36, DD = −0.37; LOS = 0.45; DHD with DD = 0.13; DHD with LOS = −0.26; DD with LOS = −0.28). Table S5 shows that the LOS-by-gender-by-region interaction was significant (F(11, 2474.63) = 1.94, p = 0.03). LOS was associated with smaller volumes in women and larger volumes in men in the CA1 (t(257.95) = −2.493, p = 0.01). CA1 volumes increased by 0.03% per year and decreased by 0.02% per year of sobriety in AUDm and AUDw, respectively.

## Discussion

Results of this study confirmed findings of smaller volumes of hippocampal subfields in association with AUD [17,22,23]. Compared to the NC group, the AUD participants exhibited significantly smaller volumes in the CA1, subiculum (with marginal significance), molecular layer, and hippocampal tail. Smaller volumes in these regions could result in abnormal neural processing, coincident with impairments of distinct mental functions. We also found significant associations between the volumes of individual subfields with age, memory, gender, and measures of drinking history.

Alterations in hippocampal subfield volumes have implications both for downstream targets and for hippocampal processing of upstream inputs. Accordingly, fewer neurons in the CA1 and subiculum may result in weaker output from the hippocampus, and smaller dendritic arborizations in the molecular layer may cause the hippocampus (particularly anterior regions) to be less effective in distinguishing between distinct contextual inputs, thereby resulting in forms of overgeneralization [54]. Both types of abnormality (input and output) could have severe consequences for emotion and motivation or sensitivity to reward. For example, the hippocampus projects to medial prefrontal areas, which are involved in conflict detection [55]. Weak or insufficiently precise contextual signals from hippocampus may bias these cortices towards greater sensitivity to emotional signals, such as those received from the amygdala. Elevated or overgeneralized processing of conflict may, in turn, bias the motivational and reward systems, particularly the nucleus accumbens, towards the use of poor coping strategies, including excessive drinking and other forms of self-medication. Abnormal inputs also can bias processing. For example, since the hippocampus receives connections from the amygdala [56], those signals may modulate the hippocampus more strongly if it has fewer neurons or smaller dendritic arborizations. This could act synergistically with weak contextual representations, thereby reinforcing their emotional valence. In other words, overgeneralized contextual representations might be more susceptible to “somatic markers” [57], as well as other forms of associative learning [54], which would increase the vulnerability to contextual signals that motivate drinking.

### CA1 and Subiculum

Both output regions of the hippocampus (CA1 and subiculum) were reduced in volume, which could be related to two cognitive corollaries: (1) an abnormal ability to distinguish a currently experienced context from other similar previously learned contexts [58, 59] and (2) higher susceptibility to emotion-driven actions [60]. That is, following a history of AUD, a given experienced context might be more likely to match a previous context in which drinking occurred.

The role of context in triggering alcohol craving is well established, and context has behavioral and treatment implications [61, 62]. Evidence suggests that social support may be helpful for circumventing specific contexts entirely [63], which would avoid the aforementioned overgeneralizing activity of an impaired CA1 and subiculum. The subiculum in particular has been implicated in the prediction of future rewards [64], perhaps due to its projections (along with projections of CA1) to the nucleus accumbens and ventral tegmental area. Likewise, the CA1 has been shown to play a role in mental processes involving envisioning the self in the future and past [41]. Therefore, the abnormalities we observed in these structures in association with AUD could negatively impact motivation through the impairment of both future processing and reward prediction.

### Molecular Layer

The molecular layer also was smaller in the AUD group. This region includes the apical dendrites of hippocampal pyramidal cells, and is traditionally viewed as an input region to the subfields of the hippocampus [65]. Smaller volumes of the molecular layer could suggest a relationship of AUD with hippocampal inputs or with local computational processes of the hippocampus (as opposed to connectivity exiting the structure) and could lead to outcomes that are similar to those resulting from smaller CA1 and subiculum volumes. In the molecular layer, entorhinal cortex inputs are thought to convey information from the current sensory context, while intra-hippocampal projections to the molecular layer are thought to represent a memory representation of the current context [66, 67]. With reduced volume of this circuitry in the molecular layer, the quality of the processing at the meeting of these two streams of input could be impacted. Therefore, it could result in a reduced ability to compare previously experienced contexts with the current experience, and might lead the hippocampus to overgeneralize the present context to previously experienced high-valence contexts. In turn, this could bias the system to addictive behaviors, i.e., similar outcomes to those discussed regarding reductions in CA1 and subiculum volumes.

### Hippocampal Tail

While the anterior hippocampus has been associated with stress, emotion, and affect [42], reductions in cells and cell processes in the posterior tail could involve many elements of spatial processing. This possibility is congruent with previous findings of alcohol-mediated alterations of spatial processing [68–70], which in rats, has been hypothesized to occur through alterations of hippocampal place cells [71]. In our AUD group, hippocampal tail volume was negatively associated with both immediate and delayed memory scores, in contrast to the positive association observed in the NC group. One possible interpretation for this finding involves regional compensation within the brain [70,72,73]. That is, for AUD participants with smaller hippocampal tail volumes, the extent of structural impairment might be sufficient to necessitate a shift to using other structures for the same function. Once this shift occurs, the new structures may provide good compensation for learning — but this shift may occur more often for the more extreme cases of hippocampal perturbation. In contrast, for AUD participants with mild hippocampal impairments, the brain may continue to rely on the impaired structures instead of shifting to an alternative compensatory region; relying on the impaired structures could result in impaired memory performance. (A metaphor that may assist in understanding is as follows: One may continue using a somewhat functioning toaster and make low quality toast; with a broken toaster one may switch to the oven broiler and make better toast.) Taken together, this explanation would fit with the observed tendency for larger hippocampal volumes to be associated with worse performance in the AUD group, and is consistent with findings that report how medial prefrontal cortex may compensate by increasing its efficiency for learning and memory after substantial hippocampal dysfunction [74].

### Influence of Participant Characteristics

While our results of smaller volumes of the CA1, subiculum, and molecular layer in AUD subjects are in agreement with past findings [17,22,23], our results did not indicate significant reductions in other regions that were reported in those studies, including CA2+3, CA4, HATA, and fimbria. However, abstinence durations for the AUD participants examined in those papers were generally shorter than in the present study (an average of 7.1 years). Therefore, results reported in the earlier studies could reflect both acute effects of alcohol consumption and long-term sequelae of chronic alcohol use. For the regions that are consistent across those studies and ours (CA1, molecular layer, and subiculum), the combined results support the view that they are vulnerable to chronic long-term AUD.

Gender differences also contribute to divergent findings across studies. As a proportion of head size, total hippocampal volumes are larger in women than in men [21, 75], a finding we support in more detail with evidence from subfield volumetric analyses (subiculum, presubiculum, CA1, and molecular layer). Although the present study and the results reported by Agartz et al. [21] and Zahr et al. [23] did not show significant interactions between gender and diagnostic group, gender differences in volumes may be elucidated further by considering quantity and duration of alcohol consumption, and LOS. In the present study, CA1 volume was related to LOS differently for men and for women. In men, CA1 volume increased with LOS, suggesting recovery over time. However, in women, the volumes continued to decline with sobriety, even after statistically accounting for age. This might indicate an AUD-related abnormality in another biological system, as AUDw may be more susceptible to liver injury and heart disease, and have been reported to display lower drinking thresholds for systemic damage [76]. Thus, a brain abnormality could continue to result in reduced health over a long period, even following abstinence, explaining the divergent gender effects of LOS. Additionally, in mice, chronic ethanol administration induces astrocyte activation in a number of hippocampal regions, including the CA1, but exclusively in female mice [77]. Despite recovery from chronic ethanol intoxication, female mice exhibited altered astrocyte neurochemistry and function. Astrocytes are involved in neural health and vascular coupling, so abnormalities could have substantial neural consequences. While further studies are needed to translate these results to human populations, this study suggests that women could be more vulnerable to life-long astrocyte inflammation-mediated brain damage, including in the hippocampus and associated regions [77].

Age, another factor that individuates the impact of AUD, was more negatively associated with hippocampal volumes in our AUD group than in our NC group for the subiculum, presubiculum, hippocampal tail, and molecular layer regions. These findings extend earlier reports based on pathology and MRI investigations, that brain regions, including the hippocampus, show greater volumetric reductions or abnormal blood flow in older than in younger individuals with AUD [19, 32]. Moreover, cognitive ramifications of an interaction of age and AUD were reported to exert compounded abnormalities in memory and visuospatial abilities [78]. In the present study, age was negatively associated with volumes of the subiculum, presubiculum, hippocampal tail, and molecular layer regions, whereas Zahr and colleagues [23] found smaller volumes in the CA2+3 subfield. While not resolved, it should be noted that there is a long-running hypothesis discussing how age may be associated with greater volume losses and functional decline in AUD groups than NC groups [79, 80].

### Limitations

In the present study, the automated subfield labeling procedure that we employed relied upon a probabilistic atlas, rather than borders defined by image contrast at high resolution. Additionally, we used a cross-sectional design, whereas a longitudinal cohort may be better able to show how the observed abnormalities were related to pre-existing risk factors for AUD or consequences of AUD. We did not consider several other factors that could influence our findings (see Oscar-Berman et al. [70] for review). For example, we did not consider additional participant characteristics (Table S6) such as family history of AUD [81] or smoking [82, 83]. As described in the Methods, although we included participants with confounding factors to increase the generalizability to the United States population, we recognize that these confounding factors could limit interpretability. For that reason, we analyzed the data from a subsample of participants without the confounding factors. All statistical group effects, including group interactions, remained significant for this unconfounded subsample. As a separate issue, the AUDw group had longer LOS and shorter DHD than the AUDm group. As described in the Methods, we analyzed subsamples of AUDm and AUDw who did not differ significantly on LOS or DHD, and significant gender effects remained significant. For simplicity, measures of education and IQ were not included in the statistical models presented. However, when the models were re-run controlling for age and education, all statistical group effects, including group interactions, remained significant.

## Conclusions

The results indicated smaller input (molecular layer) and output (CA1) volumes for the AUD group, abnormalities that could be related to distorted context processing in the AUD group. Smaller volumes also were evident for the hippocampal tail, implicating deficits in spatial processing. Memory scores were negatively associated with hippocampal tail volume in the AUD group, while a positive association was observed for the NC group, suggesting that the larger volumes were associated with better performance. This finding might further signal an AUD-related functional deficit in the hippocampal tail, and the spatial processing and memory functions performed by that region. Our observation of more extreme age-related hippocampal volume reductions in the AUD group than in the NC group, not only are congruent with the notion of synergistic negative impacts of alcohol exposure and aging; they also refine the subregional implications of the abnormality to the hippocampal tail, subiculum, presubiculum, and molecular layer. Regarding gender differences, longer LOS was associated with larger CA1 volumes in AUDm, possibly indicative of recovery of contextual processing over time; however, the smaller volumes observed in AUDw in conjunction with longer sobriety periods, might suggest impaired recovery for women. We believe that our findings not only build upon other work that highlights brain structural and functional abnormalities in the impact of AUD, they also suggest that clinicians, educators, and public health officials could benefit by approaching prevention and treatment strategies with respect to individual differences.

## Acknowledgements

This work was supported by funds from the National Institute on Alcohol Abuse and Alcoholism (NIAAA) of the National Institutes of Health US Department of Health and Human Services under Award Numbers R01AA07112 and K05AA00219; US Department of Veterans Affairs Clinical Science Research and Development grant I01CX000326; and shared instrumentation grants 1S10RR023401, 1S10RR019307, and 1S10RR023043 from the National Center for Research Resources (now National Center for Advancing Translational Sciences) at the Athinoula A. Martinos Center, Massachusetts General Hospital, and Boston University Clinical and Translational Sciences Institute (BU CTSI; 1UL1TR001430). The authors thank Zoe Gravitz, Yohan John, Steve Lehar, Riya Luhar, Nikos Makris, Diane Merritt, Greg Millington, Pooja Parikh, Jason Tourville, Maria Valmas, and Andrew Worth for assistance with recruitment, assessment, data analyses, neuroimaging, or manuscript preparation. The content is solely the responsibility of the authors and does not necessarily represent the official views of the National Institutes of Health, the U.S. Department of Veterans Affairs, or the United States Government.

## Supporting Tables and Figures

**Table S1.**
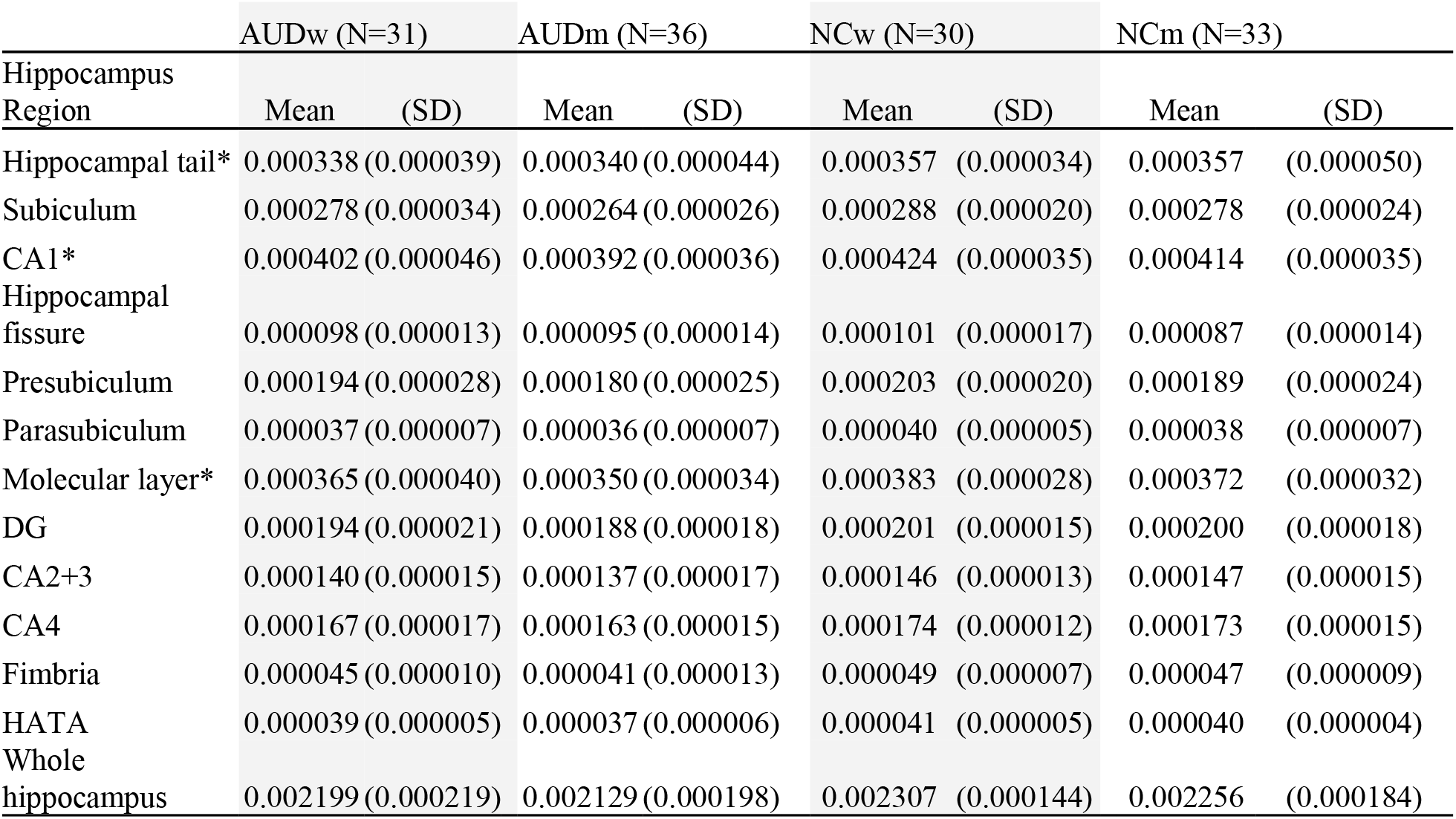
Regional volumes (proportion of estimated total intracranial volume). Means and standard deviations (SD) are provided for the hippocampal regional volumes of AUDw (N=31) and AUDm (N=36) (women and men with a history of Alcohol Use Disorder), along with NCw (N=30) and NCm (N=33) (women and men without a history of AUD). Abbreviations: CA1 through 4 = cornu ammonis 1 through 4; DG = dentate gyrus; HATA = hippocampal-amygdaloid transition area. *Indicates regions where AUD < NC, *p* < 0.0042.

**Table S2.** Analysis of variance for model including group, gender, and age. The analysis of variance obtained from the mixed model significant indicated significant group-by-region-by-age and gender-by-region-by-age interactions for volumes. Colons indicate interaction effects. Abbreviations: Sum Sq = sums of squares; Mean Sq = mean square; NumDF = numerator degrees of freedom; DenDF = denominator degrees of freedom; Pr(>F) = probability > F (i.e., *p* value).

|  | Sum Sq | Mean Sq | NumDF | DenDF | F value | Pr(>F) |
| --- | --- | --- | --- | --- | --- | --- |
| group | 1.3E-09 | 1.3E-09 | 1 | 78.97 | 4.11 | 0.05 |
| region | 1.8E-06 | 1.6E-07 | 11 | 5366.83 | 519.09 | <0.01 |
| age | 7.4E-09 | 7.4E-09 | 1 | 78.97 | 23.77 | <0.01 |
| gender | 1.3E-09 | 1.3E-09 | 1 | 78.97 | 4.18 | 0.04 |
| group:region | 6.2E-09 | 5.7E-10 | 11 | 5366.83 | 1.82 | 0.05 |
| group:age | 2.7E-09 | 2.7E-09 | 1 | 78.97 | 8.81 | <0.01 |
| region:age | 5.4E-08 | 4.9E-09 | 11 | 5366.83 | 15.59 | <0.01 |
| region:gender | 1.2E-08 | 1.1E-09 | 11 | 5366.83 | 3.41 | <0.01 |
| age:gender | 6.3E-10 | 6.3E-10 | 1 | 78.97 | 2.01 | 0.16 |
| group:gender | 2.8E-10 | 2.8E-10 | 1 | 78.97 | 0.88 | 0.35 |
| group:region:age | 1.0E-08 | 9.4E-10 | 11 | 5366.83 | 3.02 | <0.01 |
| region:age:gender | 8.6E-09 | 7.8E-10 | 11 | 5366.83 | 2.50 | <0.01 |
| group:region:gender | 2.3E-09 | 2.0E-10 | 11 | 5366.83 | 0.66 | 0.78 |
| group:age:gender | 3.9E-10 | 3.9E-10 | 1 | 78.97 | 1.25 | 0.27 |

**Table S3.** Analysis of variance for model including Immediate Memory Index. The analysis of variance obtained from the mixed model indicated a significant group-by-region-by-IMI interaction for volumes. Colons indicate interaction effects. Abbreviations: Sum Sq = sums of squares; Mean Sq = mean square; NumDF = numerator degrees of freedom; DenDF = denominator degrees of freedom; Pr(>F) = probability > F (i.e., *p* value).

|  | Sum Sq | Mean Sq | NumDF | DenDF | F value | Pr(>F) |
| --- | --- | --- | --- | --- | --- | --- |
| group | 3.5E-10 | 3.5E-10 | 1 | 75.01 | 1.12 | 0.29 |
| region | 4.9E-07 | 4.5E-08 | 11 | 5274.22 | 144.84 | <0.01 |
| IMI | 3.3E-11 | 3.3E-11 | 1 | 75.01 | 0.11 | 0.75 |
| gender | 1.3E-09 | 1.3E-09 | 1 | 75.01 | 4.05 | 0.05 |
| age | 6.7E-09 | 6.7E-09 | 1 | 75.01 | 21.82 | <0.01 |
| group:region | 6.1E-09 | 5.5E-10 | 11 | 5274.22 | 1.80 | 0.05 |
| group:IMI | 8.0E-11 | 8.0E-11 | 1 | 75.01 | 0.26 | 0.61 |
| region:IMI | 5.9E-10 | 5.4E-11 | 11 | 5274.22 | 0.17 | 1.00 |
| region:gender | 9.3E-09 | 8.4E-10 | 11 | 5274.22 | 2.73 | <0.01 |
| gender:age | 5.7E-10 | 5.7E-10 | 1 | 75.01 | 1.85 | 0.18 |
| region:age | 4.3E-08 | 3.9E-09 | 11 | 5274.22 | 12.68 | <0.01 |
| group:gender | 1.0E-11 | 1.0E-11 | 1 | 75.01 | 0.03 | 0.85 |
| group:age | 1.2E-09 | 1.2E-09 | 1 | 75.01 | 3.74 | 0.06 |
| group:region:IMI | 7.0E-09 | 6.3E-10 | 11 | 5274.22 | 2.06 | 0.02 |
| region:gender:age | 6.8E-09 | 6.2E-10 | 11 | 5274.22 | 2.01 | 0.02 |
| group:gender:age | 1.7E-11 | 1.7E-11 | 1 | 75.01 | 0.06 | 0.81 |
| group:region:gender | 1.2E-09 | 1.1E-10 | 11 | 5274.22 | 0.37 | 0.97 |

**Table S4.** Analysis of variance for model including Delayed Memory Index. The analysis of variance obtained from the mixed model indicated a significant group-by-region-by-DMI interaction for volumes. Colons indicate interaction effects. Abbreviations: DMI = Delayed Memory Index; Sum Sq = sums of squares; Mean Sq = mean square; NumDF = numerator degrees of freedom; DenDF = denominator degrees of freedom; Pr(>F) = probability > F (i.e., *p* value).

|  | Sum Sq | Mean Sq | NumDF | DenDF | F value | Pr(>F) |
| --- | --- | --- | --- | --- | --- | --- |
| group | 5.5E-10 | 5.5E-10 | 1 | 73.62 | 1.80 | 0.18 |
| region | 4.6E-07 | 4.2E-08 | 11 | 5276.69 | 136.07 | <0.01 |
| DMI | 7.5E-14 | 7.5E-14 | 1 | 73.62 | 0.00 | 0.99 |
| gender | 1.4E-09 | 1.4E-09 | 1 | 73.62 | 4.64 | 0.03 |
| age | 6.8E-09 | 6.8E-09 | 1 | 73.62 | 22.22 | <0.01 |
| group:region | 9.0E-09 | 8.2E-10 | 11 | 5276.69 | 2.67 | <0.01 |
| group:DMI | 2.2E-10 | 2.2E-10 | 1 | 73.62 | 0.71 | 0.40 |
| region:DMI | 7.4E-10 | 6.8E-11 | 11 | 5276.69 | 0.22 | 1.00 |
| region:gender | 1.1E-08 | 9.9E-10 | 11 | 5276.69 | 3.23 | <0.01 |
| gender:age | 6.9E-10 | 6.9E-10 | 1 | 73.62 | 2.27 | 0.14 |
| region:age | 4.4E-08 | 4.0E-09 | 11 | 5276.69 | 13.15 | <0.01 |
| group:gender | 1.1E-11 | 1.1E-11 | 1 | 73.62 | 0.04 | 0.85 |
| group:age | 1.1E-09 | 1.1E-09 | 1 | 73.62 | 3.63 | 0.06 |
| group:region:DMI | 1.0E-08 | 9.4E-10 | 11 | 5276.69 | 3.07 | <0.01 |
| region:gender:age | 8.4E-09 | 7.6E-10 | 11 | 5276.69 | 2.49 | <0.01 |
| group:gender:age | 1.9E-11 | 1.9E-11 | 1 | 73.62 | 0.06 | 0.80 |
| group:region:gender | 1.3E-09 | 1.2E-10 | 11 | 5276.69 | 0.38 | 0.97 |

**Table S5.** Analysis of variance for model including drinking history (DHD, DD, and LOS) of the AUD group. The analysis of variance obtained from the mixed model indicated a significant gender-by-region-by-LOS interaction for volumes, for the AUD group. Colons indicate interaction effects. Abbreviations: DHD = duration of heavy drinking; DD = daily drinks; LOS = length of sobriety; Sum Sq = sums of squares; Mean Sq = mean square; NumDF = numerator degrees of freedom; DenDF = denominator degrees of freedom; Pr(>F) = probability > F (i.e., *p* value).

|  | Sum Sq | Mean Sq | NumDF | DenDF | F value | Pr(>F) |
| --- | --- | --- | --- | --- | --- | --- |
| gender | 8.2E-13 | 8.2E-13 | 1 | 36.00 | 0.00 | 0.96 |
| region | 4.4E-07 | 4.0E-08 | 11 | 2474.63 | 129.65 | <0.01 |
| DHD | 4.1E-11 | 4.1E-11 | 1 | 36.00 | 0.13 | 0.72 |
| DD | 3.2E-10 | 3.2E-10 | 1 | 36.00 | 1.04 | 0.31 |
| LOS | 2.4E-10 | 2.4E-10 | 1 | 36.00 | 0.77 | 0.38 |
| age | 2.0E-09 | 2.0E-09 | 1 | 36.00 | 6.35 | 0.02 |
| gender:region | 3.6E-09 | 3.3E-10 | 11 | 2474.63 | 1.06 | 0.39 |
| gender:DHD | 1.3E-12 | 1.3E-12 | 1 | 36.00 | 0.00 | 0.95 |
| region:DHD | 2.2E-09 | 2.0E-10 | 11 | 2474.63 | 0.65 | 0.79 |
| gender:DD | 1.9E-11 | 1.9E-11 | 1 | 36.00 | 0.06 | 0.81 |
| region:DD | 3.7E-09 | 3.4E-10 | 11 | 2474.63 | 1.09 | 0.37 |
| gender:LOS | 7.0E-10 | 7.0E-10 | 1 | 36.00 | 2.28 | 0.14 |
| region:LOS | 7.3E-09 | 6.7E-10 | 11 | 2474.63 | 2.15 | 0.01 |
| gender:age | 1.5E-10 | 1.5E-10 | 1 | 36.00 | 0.49 | 0.49 |
| region:age | 1.4E-08 | 1.2E-09 | 11 | 2474.63 | 3.97 | <0.01 |
| gender:region:DHD | 1.8E-09 | 1.6E-10 | 11 | 2474.63 | 0.52 | 0.89 |
| gender:region:DD | 3.8E-09 | 3.5E-10 | 11 | 2474.63 | 1.12 | 0.34 |
| gender:region:LOS | 6.6E-09 | 6.0E-10 | 11 | 2474.63 | 1.94 | 0.03 |
| gender:region:age | 4.5E-09 | 4.1E-10 | 11 | 2474.63 | 1.32 | 0.21 |

**Table S6.** Additional participant characteristics. Counts of participants are given for each level of the measures listed for AUDw (N=31) and AUDm (N=36) (women and men with a history of Alcohol Use Disorder), along with NCw (N=30) and NCm (N=33) (women and men without a history of AUD). Confounded and unconfounded subsample assignment is described in the Methods. First Degree History was indicated by participant endorsement of ‘Alcoholic’ for mother, father, sibling, or offspring. Second Degree History was indicated by participant endorsement of ‘Alcoholic’ for grandparents, aunts, uncles, or grandchildren.

| Measure | Level | AUDm | AUDw | NCm | NCw |
| --- | --- | --- | --- | --- | --- |
| Confounding | Confounded | 22 | 19 | 6 | 6 |
| Confounding | Unconfounded | 14 | 12 | 27 | 24 |
| Current Smoking | No | 22 | 19 | 29 | 28 |
| Current Smoking | Yes | 14 | 12 | 4 | 2 |
| Five Year Drug History | No | 31 | 27 | 31 | 27 |
| Five Year Drug History | Less than once per week | 4 | 2 | 1 | 2 |
| Five Year Drug History | Once per week or more | 1 | 2 | 1 | 1 |
| Lifetime Drug History | No | 9 | 11 | 26 | 23 |
| Lifetime Drug History | Less than once per week | 11 | 7 | 3 | 3 |
| Lifetime Drug History | Once per week or more | 16 | 13 | 4 | 4 |
| Mother History | Alcoholic | 8 | 9 | 2 | 6 |
| Mother History | Alcoholic and drug abuse | 2 | 3 | 0 | 2 |
| Mother History | Drug abuse | 0 | 1 | 0 | 0 |
| Mother History | Never | 6 | 5 | 7 | 7 |
| Mother History | Social drinker | 20 | 13 | 24 | 15 |
| Father History | Alcoholic | 16 | 19 | 7 | 6 |
| Father History | Alcoholic and drug abuse | 3 | 3 | 0 | 2 |
| Father History | Don't know | 3 | 1 | 2 | 3 |
| Father History | Never | 1 | 2 | 3 | 4 |
| Father History | Social drinker | 13 | 6 | 21 | 15 |
| First Degree History | No | 11 | 2 | 25 | 16 |
| First Degree History | Yes | 25 | 29 | 8 | 14 |
| Second Degree History | No | 14 | 5 | 22 | 18 |
| Second Degree History | Yes | 22 | 26 | 11 | 12 |

**Figure S1.**
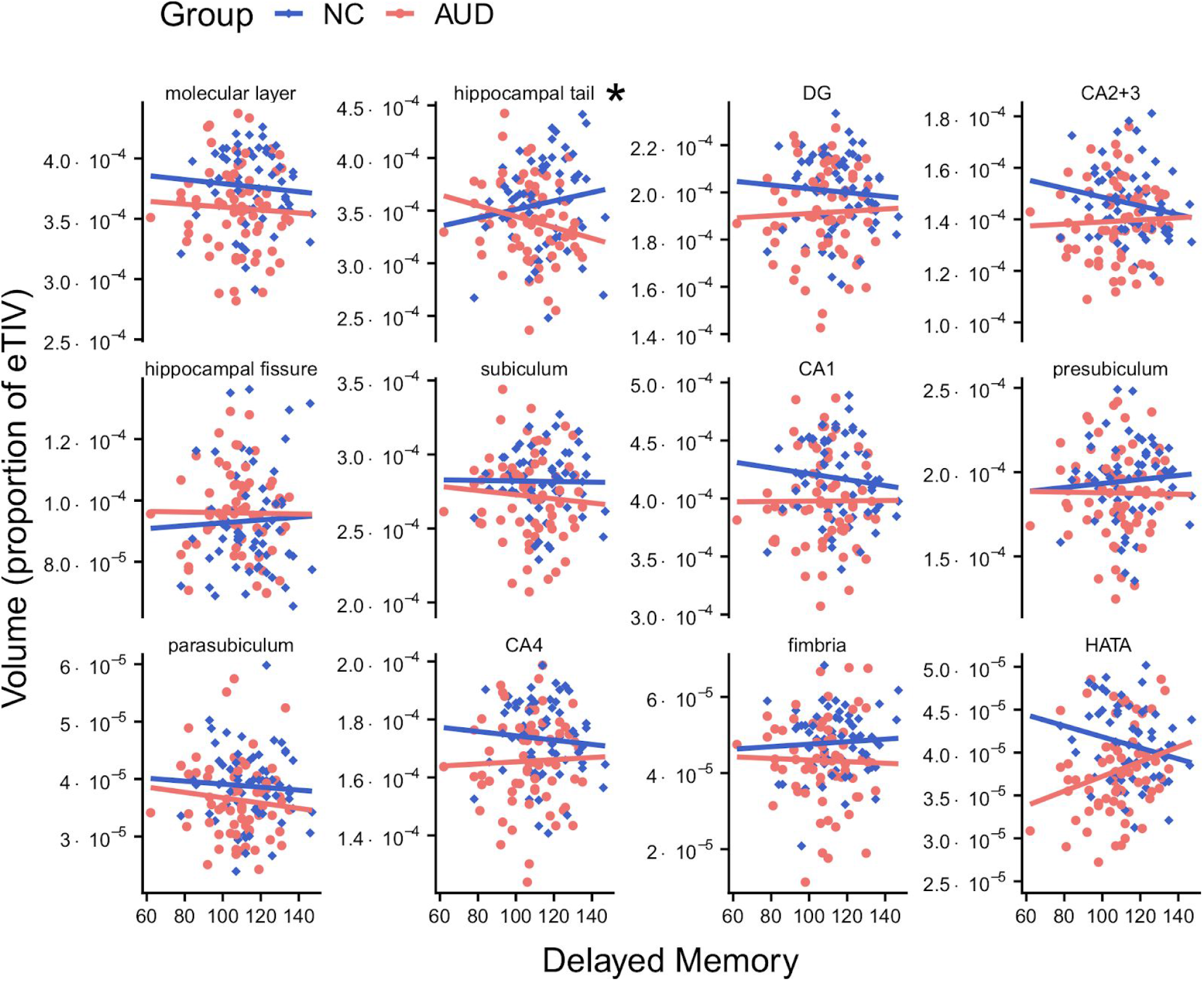
Relationships of volume with delayed memory for AUD and NC groups. As described in Figure 3, for the AUD group, Delayed Memory Index was associated with smaller hippocampal tail volumes (proportion of eTIV), while for the NC group, a positive relationship was observed. This figure shows the relationships for all 12 regions. Abbreviations: CA1 = cornu ammonis 1; CA2+3 = cornu ammonis 2 and 3; CA4 = cornu ammonis 4; DG = dentate gyrus; HATA = hippocampal-amygdaloid transition area; Sub = subiculum; eTIV = estimated total intracranial volume. *Indicates regions where *p* < 0.0042 for the interaction of group-by-Delayed Memory Index.

**Figure S2.**
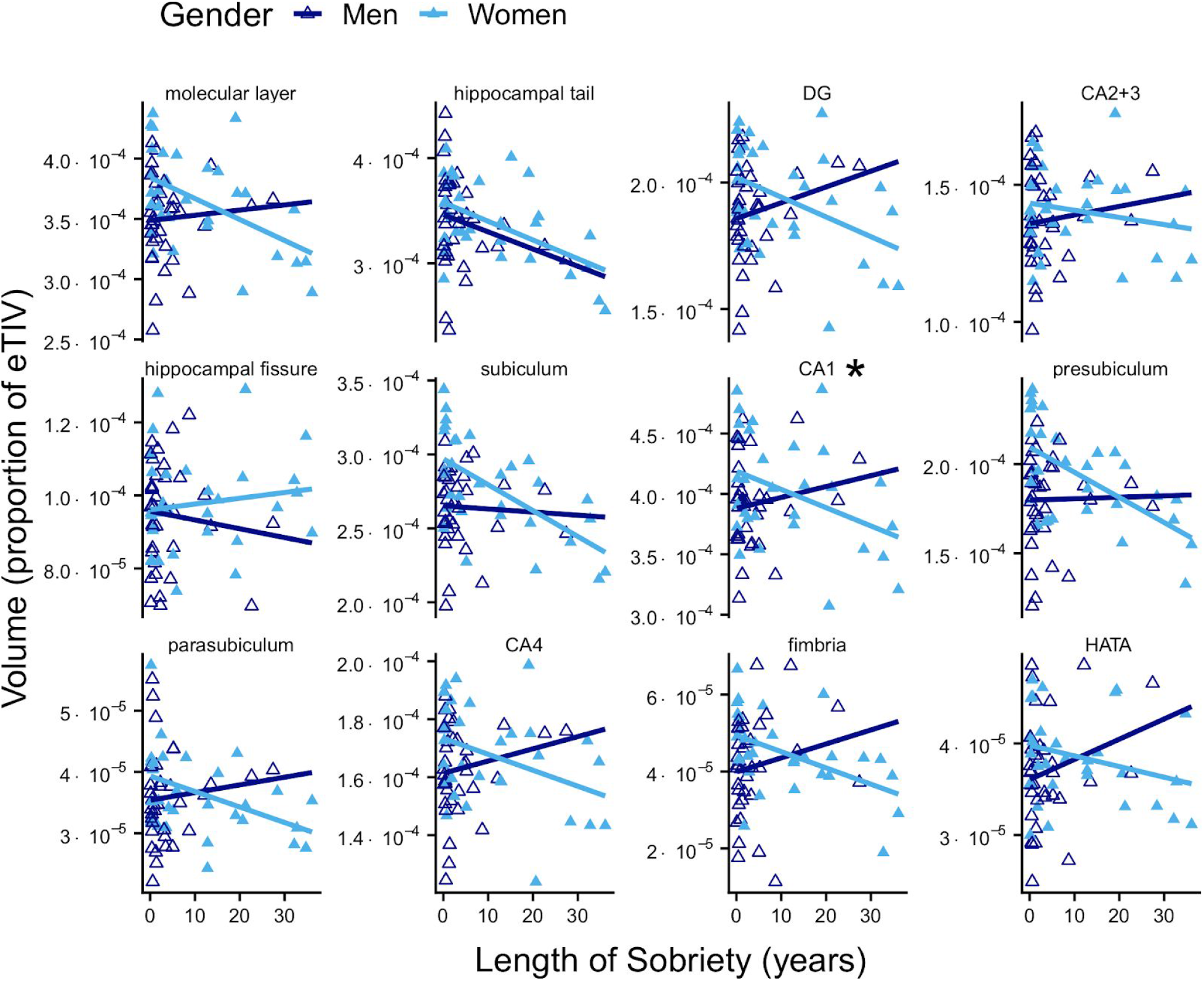
Relationships of volume with length of sobriety for AUD men and women. As described in Figure 3, for AUD men, CA1 volumes (proportion of eTIV) were positively associated with Length of Sobriety, while for AUD women, a negative relationship was observed. This figure shows the relationships for all 12 regions. Abbreviations: CA1 = cornu ammonis 1; CA2+3 = cornu ammonis 2 and 3; CA4 = cornu ammonis 4; DG = dentate gyrus; HATA = hippocampal-amygdaloid transition area; Sub = subiculum; eTIV = estimated total intracranial volume. *Indicates regions where *p* < 0.0042 for the interaction of group-by-Length of Sobriety.

## Notes

#### Summary of Updates

The manuscript was revised in response to review.

https://gitlab.com/kslays/moblab-hippocampus

## References

1. Nutt DJ, King LA, Phillips LD, Independent Scientific Committee. Drug harms in the UK: a multicriteria decision analysis. Lancet. 2010;376: 1558–1565.

2. Grant BF, Goldstein RB, Saha TD, Chou SP, Jung J, Zhang H, et al. Epidemiology of DSM-5 alcohol use disorder: Results from the National Epidemiologic Survey on Alcohol and Related Conditions III. JAMA Psychiatry. 2015;72: 757–766.

3. Staples MC, Mandyam CD. Thinking after drinking: Impaired hippocampal-dependent cognition in human alcoholics and animal models of alcohol dependence. Front Psychiatry. 2016;7: 162.

4. Fein G, Fein D. Subcortical volumes are reduced in short-term and long-term abstinent alcoholics but not those with a comorbid stimulant disorder. Neuroimage Clin. 2013;3: 47–53.

5. Wilson S, Bair JL, Thomas KM, Iacono WG. Problematic alcohol use and reduced hippocampal volume: a meta-analytic review. Psychol Med. 2017;47: 2288–2301.

6. Hommer D, Momenan R, Kaiser E, Rawlings R. Evidence for a gender-related effect of alcoholism on brain volumes. Am J Psychiatry. 2001;158: 198–204.

7. Ruiz SM, Oscar-Berman M. Closing the gender gap: The case for gender-specific alcoholism research. J Alcohol Drug Depend. 2013;1: 1–3.

8. Sawyer KS, Maleki N, Papadimitriou G, Makris N, Oscar-Berman M, Harris GJ. Cerebral white matter sex dimorphism in alcoholism: a diffusion tensor imaging study. Neuropsychopharmacology. 2018;43: 1876–1883.

9. Sawyer KS, Oscar-Berman M, Barthelemy OJ, Papadimitriou GM, Harris GJ, Makris N. Gender dimorphism of brain reward system volumes in alcoholism. Psychiatry Res. 2017;263: 15–25.

10. Mosher Ruiz S, Oscar-Berman M, Kemppainen MI, Valmas MM, Sawyer KS. Associations between personality and drinking motives among abstinent adult alcoholic men and women. Alcohol Alcohol. 2017;52: 496–505.

11. Oscar-Berman M, Ruiz SM, Marinkovic K, Valmas MM, Harris GJ, Sawyer KS. Brain responsivity to emotional faces differs in alcoholic men and women. bioRxiv. 2019. doi:10.1101/496166

12. Rivas-Grajales AM, Sawyer KS, Karmacharya S, Papadimitriou G, Camprodon JA, Harris GJ, et al. Sexually dimorphic structural abnormalities in major connections of the medial forebrain bundle in alcoholism. NeuroImage: Clinical. 2018;19: 98–105.

13. Sawyer KS, Oscar-Berman M, Mosher Ruiz S, Gálvez DA, Makris N, Harris GJ, et al. Associations between cerebellar subregional morphometry and alcoholism history in men and women. Alcohol Clin Exp Res. 2016;40: 1262–1272.

14. Seitz J, Sawyer KS, Papadimitriou G, Oscar-Berman M, Ng I, Kubicki A, et al. Alcoholism and sexual dimorphism in the middle longitudinal fascicle: a pilot study. Brain Imaging Behav. 2017;11: 1006–1017.

15. Sawyer KS, Maleki N, Urban T, Marinkovic K, Karson S, Ruiz SM, et al. Alcoholism gender differences in brain responsivity to emotional stimuli. Elife. 2019;8: e41723.

16. Beresford TP, Arciniegas DB, Alfers J, Clapp L, Martin B, Du Y, et al. Hippocampus volume loss due to chronic heavy drinking. Alcohol Clin Exp Res. 2006;30: 1866–1870.

17. Lee J, Im S-J, Lee S-G, Stadlin A, Son J-W, Shin C-J, et al. Volume of hippocampal subfields in patients with alcohol dependence. Psychiatry Res Neuroimaging. 2016;258: 16–22.

18. Chanraud S, Martelli C, Delain F, Kostogianni N, Douaud G, Aubin H-J, et al. Brain morphometry and cognitive performance in detoxified alcohol-dependents with preserved psychosocial functioning. Neuropsychopharmacology. 2007;32: 429–438.

19. Laakso MP, Vaurio O, Savolainen L, Repo E, Soininen H, Aronen HJ, et al. A volumetric MRI study of the hippocampus in type 1 and 2 alcoholism. Behav Brain Res. 2000;109: 177–186.

20. Ozsoy S, Durak AC, Esel E. Hippocampal volumes and cognitive functions in adult alcoholic patients with adolescent-onset. Alcohol. 2013;47: 9–14.

21. Agartz I, Momenan R, Rawlings RR, Kerich MJ, Hommer DW. Hippocampal volume in patients with alcohol dependence. Arch Gen Psychiatry. 1999;56: 356–363.

22. Kühn S, Charlet K, Schubert F, Kiefer F, Zimmermann P, Heinz A, et al. Plasticity of hippocampal subfield volume cornu ammonis 2+3 over the course of withdrawal in patients with alcohol dependence. JAMA Psychiatry. 2014;71: 806–811.

23. Zahr NM, Pohl KM, Saranathan M, Sullivan EV, Pfefferbaum A. Hippocampal subfield CA2+3 exhibits accelerated aging in Alcohol Use Disorder: A preliminary study. Neuroimage Clin. 2019;22: 101764.

24. Clark RE, Zola SM, Squire LR. Impaired recognition memory in rats after damage to the hippocampus. J Neurosci. 2000;20: 8853–8860.

25. Squire LR, Zola SM. Amnesia, memory and brain systems. Philos Trans R Soc Lond B Biol Sci. 1997;352: 1663–1673.

26. Barber R, McKeith IG, Ballard C, Gholkar A, O’Brien JT. A comparison of medial and lateral temporal lobe atrophy in dementia with Lewy bodies and Alzheimer’s disease: magnetic resonance imaging volumetric study. Dement Geriatr Cogn Disord. 2001;12: 198–205.

27. Mungas D, Reed BR, Jagust WJ, DeCarli C, Mack WJ, Kramer JH, et al. Volumetric MRI predicts rate of cognitive decline related to AD and cerebrovascular disease. Neurology. 2002;59: 867–873.

28. Petersen RC, Jack CR Jr, Xu YC, Waring SC, O’Brien PC, Smith GE, et al. Memory and MRI-based hippocampal volumes in aging and AD. Neurology. 2000;54: 581–587.

29. Kopelman MD, Lasserson D, Kingsley D, Bello F, Rush C, Stanhope N, et al. Structural MRI volumetric analysis in patients with organic amnesia, 2: correlations with anterograde memory and executive tests in 40 patients. J Neurol Neurosurg Psychiatry. 2001;71: 23–28.

30. Van Petten C. Relationship between hippocampal volume and memory ability in healthy individuals across the lifespan: review and meta-analysis. Neuropsychologia. 2004;42: 1394–1413.

31. Reitan RM. Validity of the Trail Making Test as an indicator of organic brain damage. Percept Mot Skills. 1958. Available: https://journals.sagepub.com/doi/pdf/10.2466/pms.1958.8.3.271

32. Sullivan EV, Marsh L, Mathalon DH, Lim KO, Pfefferbaum A. Anterior hippocampal volume deficits in nonamnesic, aging chronic alcoholics. Alcohol Clin Exp Res. 1995;19: 110–122.

33. Iglesias JE, Augustinack JC, Nguyen K, Player CM, Player A, Wright M, et al. A computational atlas of the hippocampal formation using ex vivo, ultra-high resolution MRI: Application to adaptive segmentation of in vivo MRI. Neuroimage. 2015;115: 117–137.

34. Duvernoy HM. The Human Hippocampus: An Atlas of Applied Anatomy. J.F. Bergmann; 1988.

35. Levy WB. A Computational Approach to Hippocampal Function. In: Hawkins RD, Bower GH, editors. Psychology of Learning and Motivation. NY: Academic Press; 1989. pp. 243–305.

36. Siekmeier PJ, Hasselmo ME, Howard MW, Coyle J. Modeling of context-dependent retrieval in hippocampal region CA1: implications for cognitive function in schizophrenia. Schizophr Res. 2007;89: 177–190.

37. Aimone JB, Deng W, Gage FH. Resolving new memories: a critical look at the dentate gyrus, adult neurogenesis, and pattern separation. Neuron. 2011;70: 589–596.

38. Eldridge LL, Engel SA, Zeineh MM, Bookheimer SY, Knowlton BJ. A dissociation of encoding and retrieval processes in the human hippocampus. J Neurosci. 2005;25: 3280–3286.

39. Mueller SG, Chao LL, Berman B, Weiner MW. Evidence for functional specialization of hippocampal subfields detected by MR subfield volumetry on high resolution images at 4 T. Neuroimage. 2011;56: 851–857.

40. Newmark RE, Schon K, Ross RS, Stern CE. Contributions of the hippocampal subfields and entorhinal cortex to disambiguation during working memory. Hippocampus. 2013;23: 467–475.

41. Bartsch T, Döhring J, Rohr A, Jansen O, Deuschl G. CA1 neurons in the human hippocampus are critical for autobiographical memory, mental time travel, and autonoetic consciousness. Proc Natl Acad Sci U S A. 2011;108: 17562–17567.

42. Fanselow MS, Dong H-W. Are the dorsal and ventral hippocampus functionally distinct structures? Neuron. 2010;65: 7–19.

43. Schultz C, Engelhardt M. Anatomy of the hippocampal formation. Front Neurol Neurosci. 2014;34: 6–17.

44. Hamilton M. A rating scale for depression. J Neurol Neurosurg Psychiatry. 1960;23: 56–62.

45. Robins LN, Cottler LB, Bucholz KK, Compton WM, North CS, Rourke K. Computerized diagnostic interview schedule for the DSM-IV (C DIS-IV). NIMH/University of Florida. 2000.

46. Wechsler D. WAIS-III, Wechsler Adult Intelligence Scale, Third Edition: WMS-III, Wechsler Memory Scale, Third Edition : Technical Manual. San Antonio, TX: Psychological Corporation; 1997. p. 426.

47. Cahalan D, Cisin IH, Crossley HM. American drinking practices: A national study of drinking behavior and attitudes. Monographs of the Rutgers Center of Alcohol Studies. 1969;6: 260.

48. Dawson DA, Archer L. Gender differences in alcohol consumption: effects of measurement. Br J Addict. 1992;87: 119–123.

49. Fung YL, Ng KET, Vogrin SJ, Meade C, Ngo M, Collins SJ, et al. Comparative utility of manual versus automated segmentation of hippocampus and entorhinal cortex volumes in a memory clinic sample. J Alzheimers Dis. 2019;68: 159–171.

50. Buckner RL, Head D, Parker J, Fotenos AF, Marcus D, Morris JC, et al. A unified approach for morphometric and functional data analysis in young, old, and demented adults using automated atlas-based head size normalization: reliability and validation against manual measurement of total intracranial volume. Neuroimage. 2004;23: 724–738.

51. R Core Team. R: A language and environment for statistical computing. Vienna, Austria; 2016. Available: https://www.R-project.org

52. Bates D, Mächler M, Bolker B, Walker S. Fitting linear mixed-effects models using lme4. Journal of Statistical Software, Articles. 2015;67: 1–48.

53. Allen M, Poggiali D, Whitaker K, Marshall TR, Kievit RA. Raincloud plots: a multi-platform tool for robust data visualization. Wellcome Open Res. 2019;4: 63.

54. Berkers R, Klumpers F, Fernández G. Medial prefrontal–hippocampal connectivity during emotional memory encoding predicts individual differences in the loss of associative memory specificity. Neurobiol Learn Mem. 2016;134: 44–54.

55. Thierry AM, Gioanni Y, Dégénétais E, Glowinski J. Hippocampo-prefrontal cortex pathway: anatomical and electrophysiological characteristics. Hippocampus. 2000;10: 411–419.

56. Wang J, Barbas H. Specificity of primate amygdalar pathways to hippocampus. J Neurosci. 2018;38: 10019–10041.

57. Foster PS, Hubbard T, Campbell RW, Poole J, Pridmore M, Bell C, et al. Spreading activation in emotional memory networks and the cumulative effects of somatic markers. Brain Inform. 2017;4: 85–93.

58. Smith DM, Bulkin DA. The form and function of hippocampal context representations. Neurosci Biobehav Rev. 2014;40: 52–61.

59. Barrientos SA, Tiznado V. Hippocampal CA1 subregion as a context decoder. J Neurosci. 2016;36: 6602–6604.

60. Bergado JA, Lucas M, Richter-Levin G. Emotional tagging--a simple hypothesis in a complex reality. Prog Neurobiol. 2011;94: 64–76.

61. Crombag HS, Bossert JM, Koya E, Shaham Y. Context-induced relapse to drug seeking: a review. Philos Trans R Soc Lond B Biol Sci. 2008;363: 3233–3243.

62. Zironi I, Burattini C, Aicardi G, Janak PH. Context is a trigger for relapse to alcohol. Behav Brain Res. 2006;167: 150–155.

63. Beattie MC, Longabaugh R. General and alcohol-specific social support following treatment. Addict Behav. 1999;24: 593–606.

64. Vorel SR, Liu X, Hayes RJ, Spector JA, Gardner EL. Relapse to cocaine-seeking after hippocampal theta burst stimulation. Science. 2001. pp. 1175–1178.

65. Van Strien NM, Cappaert NLM, Witter MP. The anatomy of memory: an interactive overview of the parahippocampal-hippocampal network. Nat Rev Neurosci. 2009;10: 272–282.

66. Vago DR, Kesner RP. Disruption of the direct perforant path input to the CA1 subregion of the dorsal hippocampus interferes with spatial working memory and novelty detection. Behav Brain Res. 2008;189: 273–283.

67. Hasselmo ME. The role of hippocampal regions CA3 and CA1 in matching entorhinal input with retrieval of associations between objects and context: theoretical comment on Lee et al. (2005). Behavioral neuroscience. 2005. pp. 342–345.

68. Fein G, Torres J, Price LJ, Di Sclafani V. Cognitive performance in long-term abstinent alcoholic individuals. Alcohol Clin Exp Res. 2006;30: 1538–1544.

69. Kopera M, Wojnar M, Brower K, Glass J, Nowosad I, Gmaj B, et al. Cognitive functions in abstinent alcohol-dependent patients. Alcohol. 2012;46: 665–671.

70. Oscar-Berman M, Valmas MM, Sawyer KS, Ruiz SM, Luhar RB, Gravitz ZR. Profiles of impaired, spared, and recovered neuropsychologic processes in alcoholism. Handb Clin Neurol. 2014;125: 183–210.

71. Matthews DB, Simson PE, Best PJ. Ethanol alters spatial processing of hippocampal place cells: a mechanism for impaired navigation when intoxicated. Alcohol Clin Exp Res. 1996;20: 404–407.

72. Chanraud S, Sullivan EV. Compensatory recruitment of neural resources in chronic alcoholism. Handb Clin Neurol. 2014;125: 369–380.

73. Chanraud S, Pitel A-L, Müller-Oehring EM, Pfefferbaum A, Sullivan EV. Remapping the brain to compensate for impairment in recovering alcoholics. Cereb Cortex. 2013;23: 97–104.

74. Krasne FB, Fanselow MS, Zelikowsky M. Design of a neurally plausible model of fear learning. Front Behav Neurosci. 2011;5: 41.

75. Perlaki G, Orsi G, Plozer E, Altbacker A, Darnai G, Nagy SA, et al. Are there any gender differences in the hippocampus volume after head-size correction? A volumetric and voxel-based morphometric study. Neurosci Lett. 2014;570: 119–123.

76. Vatsalya V, Liaquat HB, Ghosh K, Mokshagundam SP, McClain CJ. A review on the sex differences in organ and system pathology with alcohol drinking. Curr Drug Abuse Rev. 2016;9: 87–92.

77. Wilhelm CJ, Hashimoto JG, Roberts ML, Bloom SH, Andrew MR, Wiren KM. Astrocyte Dysfunction Induced by Alcohol in Females but Not Males. Brain Pathology. 2016. pp. 433–451. doi:10.1111/bpa.12276

78. Pfefferbaum A, Adalsteinsson E, Sullivan E V. Dysmorphology and microstructural degradation of the corpus callosum: Interaction of age and alcoholism. Neurobiol Aging. 2006;27: 994–1009.

79. Pfefferbaum A, Lim KO, Zipursky RB, Mathalon DH, Rosenbloom MJ, Lane B, et al. Brain gray and white matter volume loss accelerates with aging in chronic alcoholics: a quantitative MRI study. Alcohol Clin Exp Res. 1992;16: 1078–1089.

80. Oscar-Berman M, Marinković K. Alcohol: effects on neurobehavioral functions and the brain. Neuropsychol Rev. 2007;17: 239–257.

81. Cardenas VA, Studholme C, Meyerhoff DJ, Song E, Weiner MW. Chronic active heavy drinking and family history of problem drinking modulate regional brain tissue volumes. Psychiatry Research: Neuroimaging. 2005;138: 115–130.

82. Durazzo TC, Pennington DL, Schmidt TP, Mon A, Abé C, Meyerhoff DJ. Neurocognition in 1-month-abstinent treatment-seeking alcohol-dependent individuals: interactive effects of age and chronic cigarette smoking. Alcohol Clin Exp Res. 2013;37: 1794–1803.

83. Luhar RB, Sawyer KS, Gravitz Z, Ruiz SM, Oscar-Berman M. Brain volumes and neuropsychological performance are related to current smoking and alcoholism history. Neuropsychiatr Dis Treat. 2013;9: 1767–1784.

